# Atypical N-glycosylation of SARS-CoV-2 impairs the efficient binding of Spike-RBM to the human-host receptor hACE2

**DOI:** 10.1101/2021.04.09.439154

**Authors:** Gustavo Gámez, Juan A. Hermoso, César Carrasco-López, Alejandro Gómez-Mejia, Carlos E. Muskus, Sven Hammerschmidt

## Abstract

SARS-CoV-2 internalization by human host cells relies on the molecular binding of its spike glycoprotein (SGP) to the angiotensin-converting-enzyme-2 (hACE2) receptor. It remains unknown whether atypical N-glycosylation of SGP modulates SARS-CoV-2 tropism for infections. Here, we address this question through an extensive bioinformatics analysis of publicly available structural and genetic data. We identified two atypical sequons (sequences of N-glycosylation: NGV 481-483 and NGV 501-503), strategically located on the receptor-binding motif (RBM) of SGP and facing the hACE2 receptor. Interestingly, the cryo-electron microscopy structure of trimeric SGP in complex with potent-neutralizing antibodies from convalescent patients revealed covalently-linked N-glycans in NGV 481-483 atypical sequons. Furthermore, NGV 501-503 atypical sequon involves the asparagine-501 residue, whose highly-transmissible mutation N501Y is present in circulating variants of major concerns and affects the SGP-hACE2 binding-interface through the well-known *hotspot-353*. These findings suggest that atypical SGP post-translational modifications modulate the SGP-hACE2 binding-affinity affecting consequently SARS-CoV-2 transmission and pathogenesis.

## Introduction

The Severe Acute Respiratory Syndrome Coronavirus-2 (SARS-CoV-2) is the etiological agent for the coronavirus disease 2019 (COVID-19) pandemic, which is producing hundreds of millions of infected individuals and a disproportionate death toll worldwide^1,2^. SARS-CoV-2 is particularly aggressive and lethal for the elderly and patients with comorbidities, while remaining in most cases asymptomatic among children, adolescents and young adults^3,4^. Although an overwhelming research effort is ongoing to understand COVID-19 epidemiology, the molecular mechanisms explaining SARS-CoV-2 differential infections and severities among individuals, especially the etiology of its age-dependent risk profile, are still elusive. Here, through bioinformatics and structural analysis of publicly available data, we uncovered the likely correlation between the presence of N-glycans attached to atypical sequences of N-glycosylation (NGV-sequons) in the spike glycoprotein (SGP), and their potential to modulate SARS-CoV-2 tropism for infections.

The ability of SARS-CoV-2 to infect human host cells relies on the molecular interaction between its surface-exposed SGP and the dimerized receptor angiotensin-converting enzyme-2 (hACE2)^5–7^. SGP is a homo-trimer of approximately 440 kDa composed of modular protomers^16^. Each monomer of 1273 aa comprises a receptor-binding motif (RBM), inserted in a receptor-binding domain (RBD) that specifically recognizes *binding-hotspots* on hACE2 including the well-characterized *hotspot 31* and *hotspot 353*^17–20^. SARS-CoV-2 RBM targets its human receptor by fully-exposing a compact loop to the N-terminal helix of hACE2^19,20^. These structural traits of the SARS-CoV-2 RBM contributes to a higher hACE2-binding affinity, when compared to the SARS-CoV-1 RBM^19,20^. Both coronavirus SGPs and human hACE2 are glycosylated macromolecules, whose binding affinity and selectivity depend on their molecular structures and, the quantity, quality, and distribution of the oligosaccharide chains (glycans) they expose on their surfaces^21,22^.

The N-glycosylation of coronaviral SGPs by infected human host cells is a dynamic post-translational process. This process is essential for physiological functions and contributes to pathophysiological states^23^. In this regards, SGP shielding with human-derived N-glycans play a crucial role in infectivity, antibody recognition, and immune evasion for pathogenic coronaviruses, as is the case for HIV, Influenza and Lassa virus^8^. However, in order to ensure a more efficient protein-protein binding with their human host receptors, coronaviruses exhibit a very limited number of typical glycans in the SPG protein^8,9^. Considering the well-known N-glycosylation consensus (NXT and NXS, where X ≠ P), the SARS-CoV-2 *spike* gene encodes 22 canonical N-glycan sequons^9,10,12,24,25^. Thus, its SGP homo-trimer exhibits 66 N-glycosylation sites allowing a surface shielding of approximately 40% with a content of 28% of oligo-mannose-type glycans^9,11,12^. Nevertheless, none of these 22 typical N-glycosylation sites are directly positioned on the SPG-RBM^10–13^.

To our knowledge, both the identification and validation of atypical N-glycan motifs are not straightforward. They are experimentally identified in glycoproteins on the basis of the asparagine (N) deamidation, after N-glycosidase F (PNGase F) treatment, using mass spectrometry-based glycoproteomics^14^. However, this method can lead to false positives as deamidation can also be induced during sample preparation^14^. Recent site-specific profiling studies of glycoproteins from human sera and from the ovarian cancer cell line (OVCAR-3) overcame this technical limitation and identified two atypical glyco-site sequons with the NXC and NXV motifs^14,15,26,27^. In consequence, validated evidences of atypical N-glycan occupancy have been reported for major human serum glycoproteins such as von Willebrand Factor, CD69, serotransferrin, factor XI, albumin, and α−1B-glycoprotein^15,26,27^. Furthermore, these atypical N-glycosylations have a widespread presence in the proteome of human individuals^15,26,27^. Nonetheless, in the context of the COVID-19 pandemic, it remains elusive whether these atypical N-glycosylation sequons are also encoded in the genome of the original Wuhan-Hu-1 (the novel coronavirus), and how they could potentially contribute to the virus tropism of the SARS-CoV-2 variants of major concern.

In this study we uncovered evidences for the existence of N-glycans attached to the atypical NGV-motifs in key structural locations of the surface-exposed spike-glycoprotein of SARS-CoV-2. Our findings strongly support the potential N-glycosylation of these atypical sequons by humans in the COVID-19 pandemic. Moreover, we propose that the atypical N-glycosylation of these NGV motifs could modulate the propensity of the novel coronavirus to infect the human cells according to the host age, therefore highlighting potential targets for effective therapeutic and prophylactic interventions to control the COVID-19 pandemic.

## Results

### Atypical N-glycosylation sequons encoded in the SARS-CoV-2 reference genome

Current bioinformatics resources are limited to predict atypical sequences for N-glycosylation. Thus, we manually identified 75 genome-wide putative atypical N-glycosylation sequons (NXV and NXC) in the SARS-CoV-2 reference strain. We observed a particular abundance of these N-glycosylation motifs among protein-encoding genes such as the surface - exposed SGP and several non-exposed proteins (RdRP, nsp2, and nsp3) (Table 1). In total, we found 58 NXV-motifs (77.3%) and 17 NXC (22.7%). Six atypical NXV-sequons have the functional NGV motif. In one NXC the X corresponded to proline, which is a forbidden residue in the convention to identify typical N-glycosylation sites as it renders the sequons defective. It is worth noting that 34/58 (58.6%) of the NXV sequons are present in only four proteins: spike (12), nsp3 (11), RdRP (6), and nsp2 (5). Similarly, 12/17 (70.6%) of the NXC motifs were found to be distributed among the same four proteins: RdRP (4), spike (3), nsp2 (3), and nsp3 (2) (Table 1). These results show that the spike protein contains the highest number of putative atypical N-glycosylation sequons from the whole SARS-CoV-2 genome.

**Table 1.**
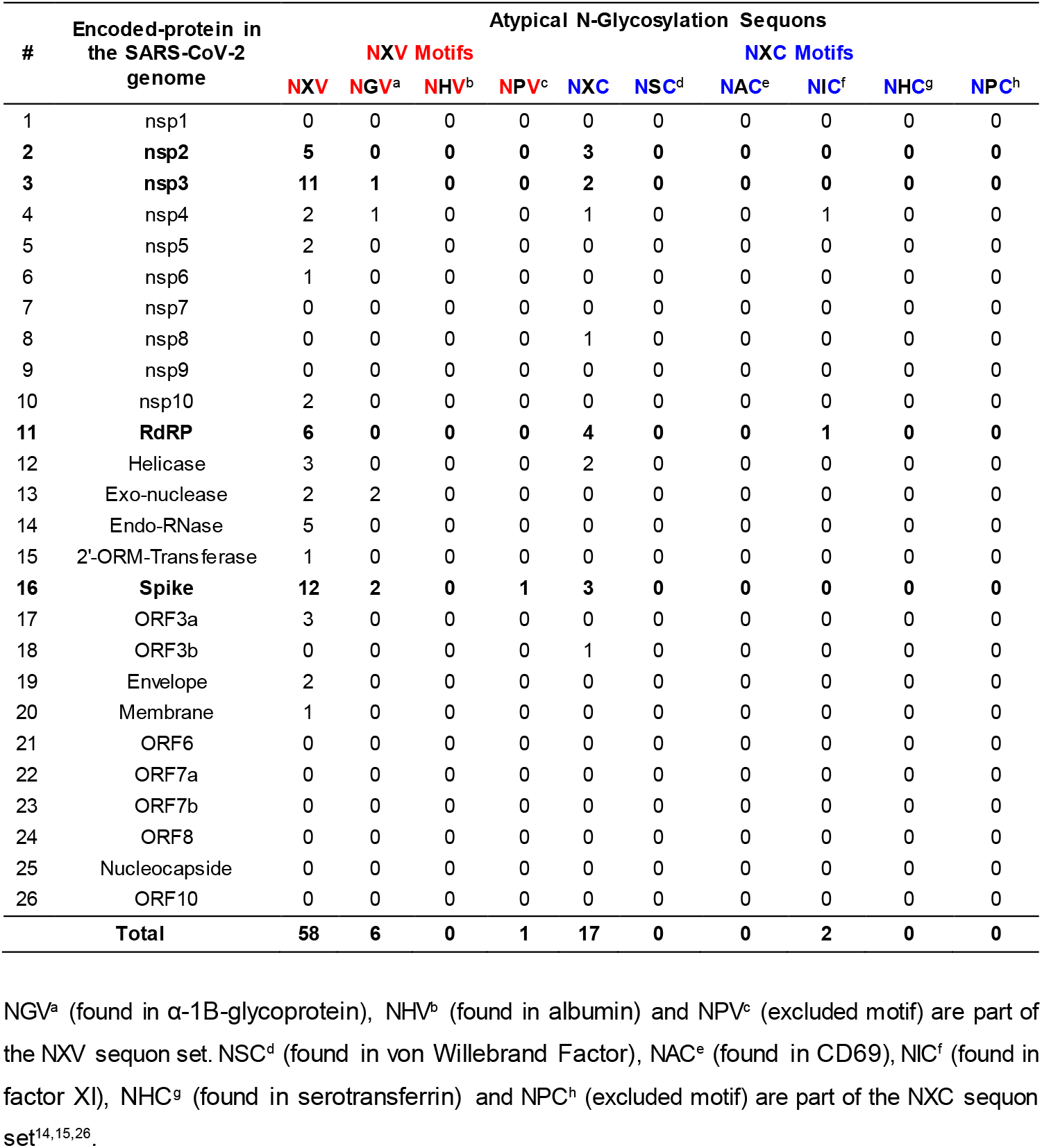
Summary of the SARS-CoV-2 genome-wide putative atypical N-glycosylation sequons identified by manual analysis and curation.

| # | Encoded-protein in the SARS-CoV-2 genome | Atypical N-Glycosylation Sequons |  |  |  |  |  |  |  |  |  |
| --- | --- | --- | --- | --- | --- | --- | --- | --- | --- | --- | --- |
|  |  | NXV Motifs |  |  |  | NXC Motifs |  |  |  |  |  |
|  |  | NXV | NGV<sup>a</sup> | NHV<sup>b</sup> | NPV<sup>c</sup> | NXC | NSC<sup>d</sup> | NAC<sup>e</sup> | NIC<sup>f</sup> | NHC<sup>g</sup> | NPC<sup>h</sup> |
| 1 | nsp1 | 0 | 0 | 0 | 0 | 0 | 0 | 0 | 0 | 0 | 0 |
| 2 | **nsp2** | **5** | **0** | **0** | **0** | **3** | **0** | **0** | **0** | **0** | **0** |
| 3 | **nsp3** | **11** | **1** | **0** | **0** | **2** | **0** | **0** | **0** | **0** | **0** |
| 4 | nsp4 | 2 | 1 | 0 | 0 | 1 | 0 | 0 | 1 | 0 | 0 |
| 5 | nsp5 | 2 | 0 | 0 | 0 | 0 | 0 | 0 | 0 | 0 | 0 |
| 6 | nsp6 | 1 | 0 | 0 | 0 | 0 | 0 | 0 | 0 | 0 | 0 |
| 7 | nsp7 | 0 | 0 | 0 | 0 | 0 | 0 | 0 | 0 | 0 | 0 |
| 8 | nsp8 | 0 | 0 | 0 | 0 | 1 | 0 | 0 | 0 | 0 | 0 |
| 9 | nsp9 | 0 | 0 | 0 | 0 | 0 | 0 | 0 | 0 | 0 | 0 |
| 10 | nsp10 | 2 | 0 | 0 | 0 | 0 | 0 | 0 | 0 | 0 | 0 |
| 11 | **RdRP** | **6** | **0** | **0** | **0** | **4** | **0** | **0** | **1** | **0** | **0** |
| 12 | Helicase | 3 | 0 | 0 | 0 | 2 | 0 | 0 | 0 | 0 | 0 |
| 13 | Exo-nuclease | 2 | 2 | 0 | 0 | 0 | 0 | 0 | 0 | 0 | 0 |
| 14 | Endo-RNase | 5 | 0 | 0 | 0 | 0 | 0 | 0 | 0 | 0 | 0 |
| 15 | 2'-ORM-Transferase | 1 | 0 | 0 | 0 | 0 | 0 | 0 | 0 | 0 | 0 |
| 16 | **Spike** | **12** | **2** | **0** | **1** | **3** | **0** | **0** | **0** | **0** | **0** |
| 17 | ORF3a | 3 | 0 | 0 | 0 | 0 | 0 | 0 | 0 | 0 | 0 |
| 18 | ORF3b | 0 | 0 | 0 | 0 | 1 | 0 | 0 | 0 | 0 | 0 |
| 19 | Envelope | 2 | 0 | 0 | 0 | 0 | 0 | 0 | 0 | 0 | 0 |
| 20 | Membrane | 1 | 0 | 0 | 0 | 0 | 0 | 0 | 0 | 0 | 0 |
| 21 | ORF6 | 0 | 0 | 0 | 0 | 0 | 0 | 0 | 0 | 0 | 0 |
| 22 | ORF7a | 0 | 0 | 0 | 0 | 0 | 0 | 0 | 0 | 0 | 0 |
| 23 | ORF7b | 0 | 0 | 0 | 0 | 0 | 0 | 0 | 0 | 0 | 0 |
| 24 | ORF8 | 0 | 0 | 0 | 0 | 0 | 0 | 0 | 0 | 0 | 0 |
| 25 | Nucleocapside | 0 | 0 | 0 | 0 | 0 | 0 | 0 | 0 | 0 | 0 |
| 26 | ORF10 | 0 | 0 | 0 | 0 | 0 | 0 | 0 | 0 | 0 | 0 |
| **Total** |  | **58** | **6** | **0** | **1** | **17** | **0** | **0** | **2** | **0** | **0** |

NGV<sup>a</sup> (found in α-1B-glycoprotein), NHV<sup>b</sup> (found in albumin) and NPV<sup>c</sup> (excluded motif) are part of the NXV sequon set. NSC<sup>d</sup> (found in von Willebrand Factor), NAC<sup>e</sup> (found in CD69), NIC<sup>f</sup> (found in factor XI), NHC<sup>g</sup> (found in serotransferrin) and NPC<sup>h</sup> (excluded motif) are part of the NXC sequon set<sup>14,15,26</sup>.

### Atypical N-glycosylation sequons located in the spike-glycoprotein (SGP)

From the 15 putative atypical N-glycosylation sequons (15.8%) present in the SGP, eleven are located in the subunit S1 and four in the subunit S2. Further, the N-terminal domain (NTD) of SGP has four atypical N-glycan motifs and the RBD bears another four sequons. In line with our expectations, two of the N-glycan sequons identified at the RBD, which exhibit the NGV-motif, are fully-exposed on its RBM domain. Moreover, both atypical N-glycan sequons (RBM positions: 481-483 and 501-503) are located in the interaction interface of SGP with the human host-receptor hACE2. In addition, the NGV 481-483 is structurally located in the middle of the ridge loop of the RBM, while the NGV 501-503 involves the critical residue N501 (Figure 1). This asparagine, N501, has been shown to interact with the hACE2-binding hotspot 353, and is a key position for major clinically concerning mutations present in several emerging SARS-CoV-2 variants^19,20^.

**Figure 1.**
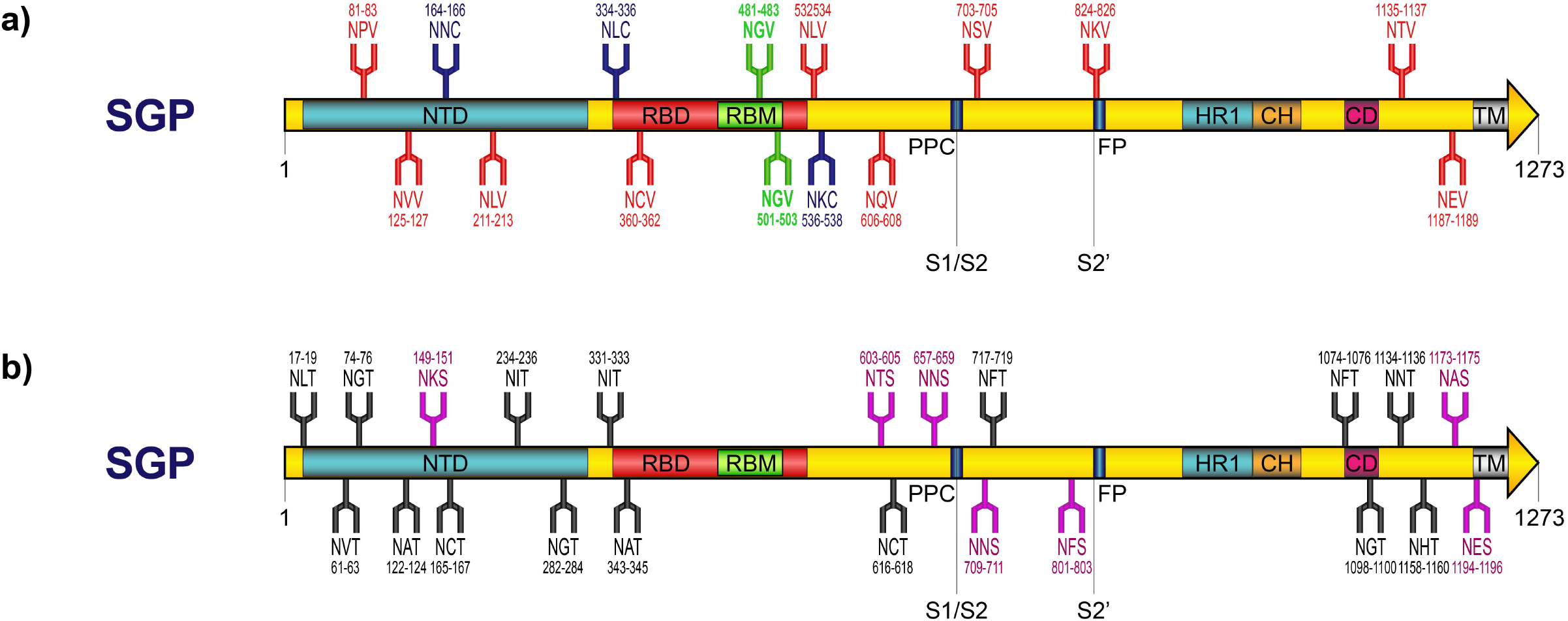
Atypical and canonical N-glycosylation sequons of the SARS-CoV-2 spike-glycoprotein. **a)** Manually-identified atypical N-glycan sites: 2 NGV (green), 10 NXV (red) and 3NXC (blue) sequons. **b)** Computer-based prediction of canonical N-glycan sites: 15 NXT (black) and 7 NXS (purple) sequons. NTD: N-terminal domain; RBD: receptor-binding domain; HR1: heptad repeat 1; CH: central helix; CD: connector domain; TM: transmembrane domain; PPC: proprotein convertase; FP: furine proteolysis. Positions in the protein are drawn to scale.

### Occupancy of atypical N-glycosylation sequons in SGP-RBM

Analysis of the three-dimensional structures of SGP and RBD available in the Protein Data Bank (PDB) (as of March 3^rd^, 2021), confirmed the presence of N-glycan disaccharides (NAG-BMA and NAG-NAG) atypically attached to the SGP-RBM ridge-loop. These disaccharides were covalently-linked to the asparagine residues of the NGV 481-483 motifs in a trimeric-SGP structure (PDB-ID: 6XEY, Figure 2A). These N-glycosylations, were observed by cryo-electron microscopy (cryo-EM) in the complex of SARS-CoV-2 spike-glycoprotein bound to an antibody fragment (Fab 2-4) identified from convalescent patients^28,29^ (Figure 2B). This structure highlights the likelihood of these atypical N-glycan sequons to be recognized and glycosylated via the human metabolism. In contrast, no atypical N-glycans were observed for the NGV 501-503 sequon among the SGP and RBD structures available in the PDB.

**Figure 2.**
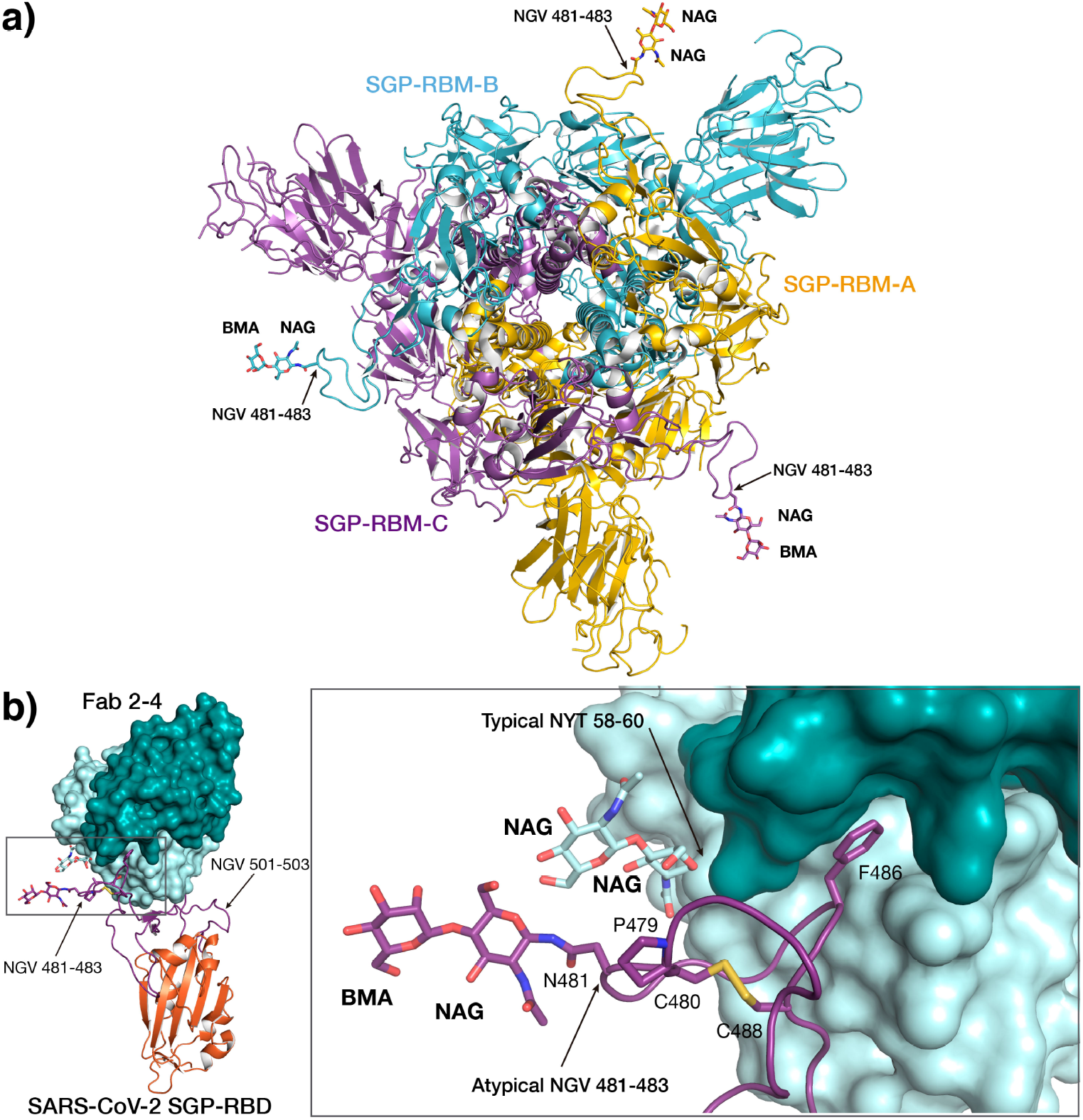
Cryo-EM structure of the SARS-CoV-2 SGP-RBM in complex with Fab 2-4 from a convalescent patient. **a)** View of the three-dimensional structure of SARS-CoV-2 SGP (PDB-ID: 6XEY) along the ternary axis. Sugars attached to the atypical N-glycosylation site NGV 481-483 for each chain are represented as capped sticks and labeled. **b)** Interaction between SARS-CoV-2 spike RBD (depicted as ribbons with RBD colored in orange and with the RBM region in violet) and Fab2-4 (depicted by its molecular surface) as observed in PDB 6XEY. Positions of the atypical N-glycosylation sites NGV 481-483 and NGV 501-503 are indicated by an arrow. Right, details of the atypical molecular interaction between N-glycans at the RBM-hACE2 binding interface. Relevant residues and sugar rings depicted as sticks and labeled. Occupancy of the atypical N-glycan NGV 481-483 site with a N-acetyl-glucosamine disaccharide (NAG-NAG/BMA) is observed in interaction with a typical but infrequent N-glycan (NYT 58-60) on the fragment antigen binding of the anti-SARS-CoV-2 antibody.

### Genetic conservation of the atypical N-glycosylation sequons across other coronaviruses

Other coronaviruses isolated from diverse species also conserve some of the atypical N-glycosylation sequons found in SARS-CoV-2. We aligned and compared the SARS-CoV-2 SGP-RBM protein sequence with the SGP-RBM of SARS-CoV-1 and other SARS-like coronaviruses isolated from pangolins and bats in China and Cambodia^30,31^. The ridge loop of the Pangolin-CoV-RBM is very similar to that of SARS-CoV-2, also conserving both atypical N-glycosylation NGV-sequons we identified in SARS-CoV-2 (Figure 3A). Nevertheless, the ridge loops of two SARS-like coronaviruses isolated from bats in Cambodia in 2010 comprise the atypical N-glycan sequon NGV 501-503, while the atypical N-glycan sequon NGV 481-483 is defective^31^. However, both atypical N-glycan motifs are defective and/or absent in the ridges from SARS-CoV-1 and the RaTG13 Bat-CoV (Figure 3A). In addition, the correspondent RBM-positions P479, C480 and C488, which are important for the conformation of the ridge loop, remain 100% conserved in all the 48 coronavirus sequences we have aligned and compared (Figure 3A).

**Figure 3.**
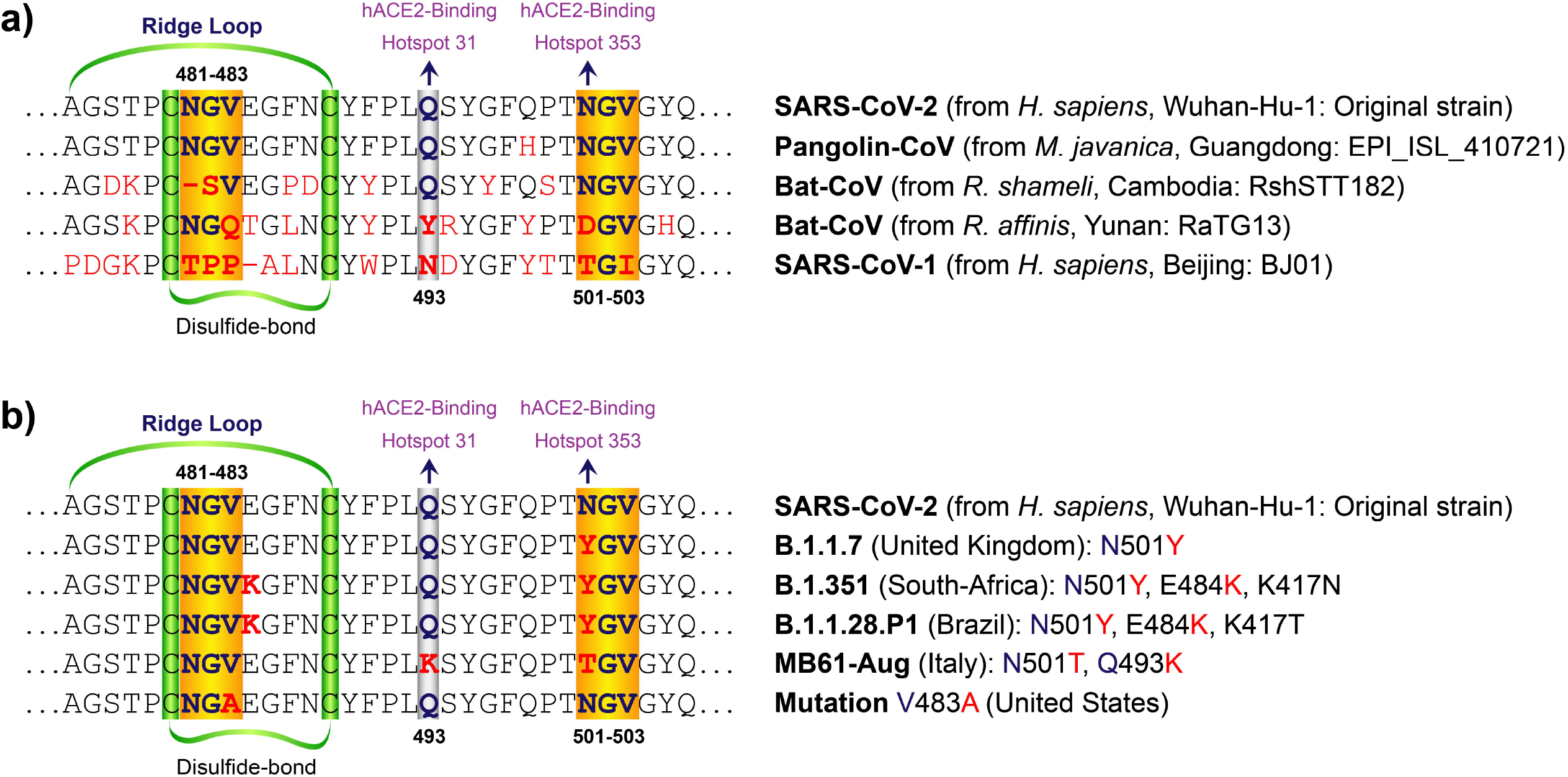
Alignment and comparison of the hACE2-binding interface of the SARS-CoV-2 SGP-RBM. **a)** Insights into the possible origin of the atypical N-glycan sequons of SARS-CoV-2 RBM. Original SARS-CoV-2 Wuhan-Hu-1 reference strain is aligned to SARS-CoV-1 (other pathogenic coronavirus), and SARS-like coronaviruses isolated from bats (RaTG13 and RshSTT182), and from a Malayan pangolin seized in Guangdong, China. **b)** Identification of SARS-CoV-2 RBM mutations (in red) affecting the atypical N-glycan sequons (light yellow boxes). A portion of the original SARS-CoV-2 Wuhan-Hu-1 strain (RefSeq: NC_045512.2) is aligned with emergent variants from United Kingdom, South-Africa, Brazil, Italy and United States.

### Genetic variability of the SGP-RBM and the atypical N-glycosylation sequons in SARS-CoV-2

We hypothesized that SGP-RBM mutations of concern affecting the RBM-hACE2 binding interface either involve the atypical N-glycosylation sequons NGV 481-483 and NGV 501-503, or are located close to them. To explore this hypothesis, we further analyzed a total of 730,744 complete genome sequences of SARS-CoV-2 (GISAID database^32^), isolated from humans (collection dates: from December 1^st^, 2019 to February 28^th^, 2021 and submission dates: from December 1^st^, 2019 to March 20^th^, 2021). We highlighted variable genetic positions, which are producing altered SARS-CoV-2 SGP-RBM variants after 15 months of pandemics. All low coverage sequences (entries with more than 5% Ns) were excluded prior to the genetic variation analysis. With the cured data, we then constructed a Variome model of the SARS-CoV-2 RBM (positions: 437-506 + 403 and 417; 72 a.a. in total) (Figure 4). We identified a total of 323 different SGP-altering changes across the 72 codons of the RBM in 259,627 reported genome sequences (Figure 4). A half of these genetic mutations (49.2%) are part of the RBM-hACE2 binding interface (positions: 475-506), with 74/323 (22.9%) being located in the ridge loop (position: 475-488) (Figure 4). Moreover, the 40 N-terminal codons of the RBM exhibited 164 variants, including the N439K (18,276x), L452R (14,553), and K417N (3,892x) Y453F (1,008x), which are also key for the RBM-hACE2 binding-interface (Figure 4). Only 143 out of the 730,744 SARS-CoV-2 genomes are found to carry seven different mutations at position N481 (Figure 4), which is the substrate of the atypical N-glycosylation sequons that are glycosylated in the trimeric-SGP structure complexed with potent neutralizing monoclonal antibodies (PDB-ID: 6XEY) (Figure 2). In contrast, the most frequent RBM mutation N501Y (165,519x) makes defective the atypical N-glycosylation sequon NGV 501-503, becoming a key RBM position for the interaction with hACE2-binding hotspot 353 (Figure 4).

**Figure 4.**
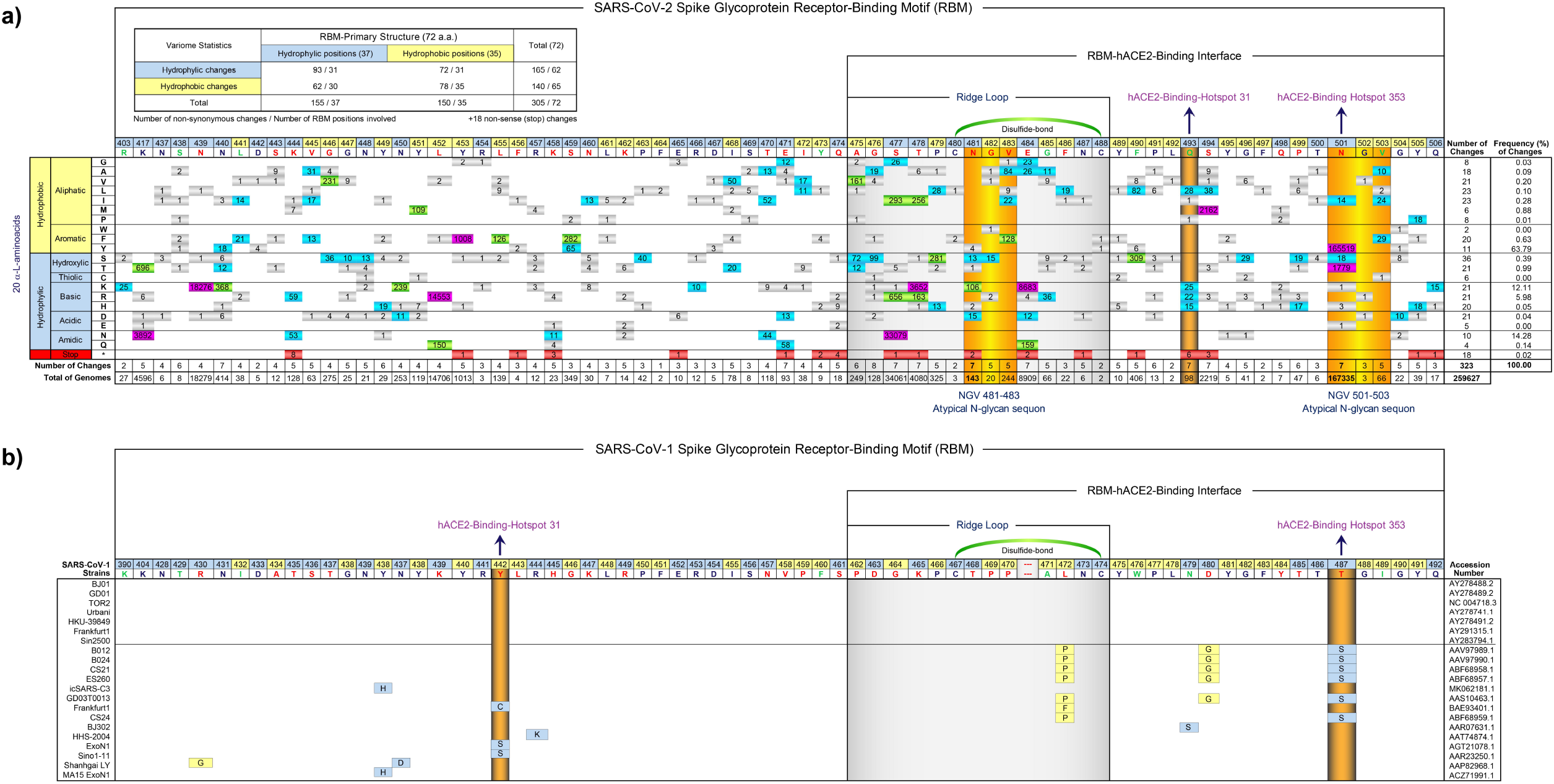
Variation analysis of the SARS-CoV-2 and SARS-CoV-1 spike-glycoproteins. The primary sequence of the receptor-binding motifs (RBM) of SARS-CoV-2 and SARS-CoV-1 are displayed and their amino-acid positions are classified according to their, hydrophobicity (light yellow) and their hydrophilicity (light blue). The 20 natural α-L-amino-acids are displayed and organized according to their physicochemical properties. For both RBM sequences, conserved amino-acid positions are highlighted in bold blue, conservative changes (different amino-acid, but similar physicochemical property) are denoted in bold green, and radical changes (different amino-acid and different physico-chemical property) are highlighted in bold red. The RBM-ridge-loops (with their disulfide-bonds) are boxed in gray for both coronaviruses, and the amino-acid positions interacting with the hACE2-binding hotspots 31 and 353 are defined by the dark yellow boxes and the arrows in the RBM-hACE2-binding interface region. The atypical N-glycosylation motifs NGV 481-483 and NGV 501-503 are highlighted in light yellow boxes. **a)** Basic statistics are presented for the Variome of the SARS-CoV-2 RBM. One-hundred and two different mutations on SGP-RBM are displayed, according to their frequency in the GISAID reported information for samples collected up to February 28th, 2021 and reported up to March 20th, 2021. Mutations with a frequency under 10 are depicted in small light gray, mutations with a frequency over 10 are depicted in small light blue, mutations with a frequency over 100 are depicted in small light green, mutations with a frequency over 1000 are depicted in small dark red and stop mutations are depicted in small dark red. N501Y is the most frequent mutation (165,519 genomes) observed up to date for the SARS-CoV-2 RBM. **b)** For the Variome of the SARS-CoV-1, 7 conserved genomic sequences were defined as references for other 14 variable genomic sequences retrieved from the NCBI (accession number are listed on the left). For SARS-CoV-1, only the amino-acid changes are presented. No mutation frequencies were estimated.

The N501Y is one of the several clinically concerning mutations located at the atypical N-glycosylation sequons. This mutation has been recently reported in the three most common SARS-CoV-2 mutant strains detected in the United Kingdom (501Y.V1), South-Africa (501Y.V2) and Brazil (501Y.V3 or P.1) (Figure 3B). Other six circulating variants have been reported for the Asn501 residue: N501T (1,779x - Italy MB61-Aug), N501S (18x), N501I (14x), N501H (3x), N501K (1x) and N501E (1x). Similarly, eight other genetic changes affecting directly the atypical N-glycan sequon NGV 501-503 have been detected up to date, and ten mutations have been identified for the NGV 481-483 (Figure 3B). The variants from South-Africa and Brazil have an additional genetic change of major concern in the ridge loop: the E484K mutation, which is located nearby the atypical N-glycan sequon NGV 481-483 (Figure 3B). The number of reported genomes including this E484K mutation can be now counted by thousands (Figure 4). Moreover, the SARS-CoV-2 variant identified in Italy presented also a non-synonymous change in the position Q493, which is important for the interaction between the SGP-RBM and the hACE2-binding hotspot 31 (Figure 3B). These findings show that mutations at atypical N-glycosylation sites are present in each of the new clinically concerning variants. It is noteworthy to mention that many of the new variants of concern harboring mutations at the atypical N-glycosylation sequons spread faster and have higher mortality rates^33,34^.

## Discussion

Despite the record achievement of several effective and safe vaccines against COVID-19 in 2020, and the current increasing vaccination rates worldwide (as of April 8^th^, 2021 a total of 669.248.795 vaccine doses have been administered)^35,36^, SARS-CoV-2 remains a significant threat for humans in 2021^1^. By re-analyzing a large collection of publicly available genetic and structural data, we established a feasible connection between atypical N-glycosylation in humans and the evolution of the COVID-19 pandemics. Atypical N-glycans may play a role in the propensity of the SARS-CoV-2 original strain and its fast-spreading genetic variants, to remain mimetically hidden among young and healthy people, while in other vulnerable individuals it causes severe infections.

While the canonical N-glycosylation sequons (NXT / NXS) of SGP have been readily predicted and identified to be well-distributed all along this 1273-aa glycoprotein^10,24^, their absence in the RBM discards their possible impact on the hACE2-binding activity of SARS-CoV-2 (Figure 1B). In contrast, the presence of atypical N-glycosylation sequons at the RBM indicated in this study could enlighten molecular explanations for the differential propensity of this aggressive coronavirus to contact target cells, internalize across the cell membrane, and to cause the acute respiratory distress syndrome (ARDS) in humans. In addition, the fact that all the clinically concerning variants have mutations at the atypical N-glycosylation sequons, highlights the importance of the N-linked sugars on these motifs in the molecular biology and epidemiology of SARS-CoV-2 pandemics. This is especially valid for the highly-conserved atypical N-glycan sequons located in the ridge-loop (NGV 481-483) (Figure 1A), whose occupancy by glycan moieties has been confirmed in a SARS-CoV-2 SGP structure (Figure 2) complexed with potent neutralizing monoclonal antibodies^29,37^. The authors of this cryo-EM trimeric-structure of the SARS-CoV-2 SGP bound to Fab 2-4 (PDB-ID: 6XEY) used the tetracycline-inducible Expi293 cell line for the high-yield expression of recombinant SARS-CoV-2 spike-glycoprotein^29,37^. Expi293^TM^ are sub-derived cells from the human embryonic kidney 293 (HEK-293) cell line, and extensively used in the production of homogeneously N-glycosylated proteins (*Thermo-Fisher Scientific*). Therefore, experimental artifacts as the origin or source of these atypical N-glycosylations of the ridge-loop can be discarded. Moreover, we also identified tight interactions between our atypical ridge-loop disaccharide sugars and the human-produced canonical N-glycan disaccharides in the antibodies (NAG-NAG in NYT 58-60 sequon of the Fab 2-4) (Figure 2). Although this observation is unusual because Fab variable regions of IgG comprise no conserved N-glycan motifs^38,39^, it is estimated that about 15% of IgG molecules can generate N-glycan sites in their Fab by somatic hyper-mutation^38,39^. It is also noteworthy to mention that these potent neutralizing antibodies were isolated from five patients infected with SARS-CoV-2, who were hospitalized with severe symptoms^29,37^. The interaction of these human-produced antibodies with the atypical N-glycosylated sites at the RBM of the SGP (produced *in vitro*) is a strong evidence for the natural relevance of the atypical NGV-sequons as potent immunogenicity epitopes of the spike protein. Thus, the interaction of Fab 2-4 with the glycan moieties attached to the NGV sequons found in the RBM, is an authentic molecular evidence that contributes to explain the role of atypical N-glycosylation sequons in the differential behavior of SARS-CoV-2 among humans in the pandemic.

SARS-CoV-2 RBM has a higher and more efficient binding affinity when compared to SARS-CoV-1, the coronavirus responsible for the epidemics in 2003. These features stem from the structural changes observed in the hACE2-binding interface of SARS-CoV-2^19,20^. Crucially, we confirmed that SARS-CoV-2 RBM has a more compact ridge-loop, which connects better with the N-terminal helix of hACE2, largely due to the six residues **N**-**G**-**V**-E-G-F (positions 481 to 486), which harbors one of the atypical N-glycosylation sequons instead of the five residues T-P-P-A-L (positions 468 to 472) present in the SARS-CoV-1 RBM ridge-loop (Figures 3 and 4). Because this SGP ridge-loop has been identified as a key genomic feature differentiating the pathogenic SARS-CoV-2, SARS-CoV-1 and MERS-CoV from less pathogenic coronaviruses^40,41^, we further aligned and compared the SGP-RBM portion corresponding to the hACE2-binding interface of SARS-CoV-2, which contains the NGV 481-483 and the NGV 501-503, with those from SARS-CoV-1 and other zoonotic coronaviruses^42^. We confirmed the presence of the atypical N-glycosylation motifs in two SARS-like CoVs isolated from bats in Cambodia and from pangolins seized in Guangdong, China, which bear at least one (NGV 501-503) or both (NGV 481-483 and NGV 501-503) sequons, respectively (Figure 3A). Conversely, SARS-CoV-1 and RaTG13 Bat-CoV (another SARS-like coronavirus isolated from bats) exhibit defective atypical N-glycan sequons in their RBM, or are absent, due to non-synonymous changes or deletions at the ridge-loop (Figure 3A). In fact, the ridge-loop and its NGV 481-483 are also absent in MERS-CoV and other coronaviruses from bats, which likely affect their hACE2 receptor binding capacity^40,41^. These observations further support the hypothesis of pangolins and bats as the natural reservoir or source of SARS-CoV-2^5,17,40^, and highpoint the importance of the atypical N-glycan sequons in SARS-CoV-2.

Our time-lapsed variome analysis of the 70-aa SGP-RBM (plus the critical positions R403 and K417 of the RBD) reveal that NGV 481-483 still remains highly conserved after one year of pandemic, while NGV 501-503 has been widely affected by concerning mutations, further enhancing our hypothesis of a determinant role of the human atypical N-glycosylation process in the COVID-19 pandemic. We observed 17 infrequent genetic changes for the atypical N-glycosylated sequon NGV 481-483, accounting for a set of just 143 out of 730,744 reported genome sequences (Figure 4). This would suggest an important functional role of the NGV motif in the original SARS-CoV-2 spike-glycoprotein, which is still maintained after genome replication as an adaptive advantage in the new SARS-CoV-2 mutants. Thus, the atypical N-glycan sequon NGV 481-483 seems to be intolerant of those loss-of-function variants, which is in agreement with the mutation-refractory behavior observed for its neighbor positions C480 and C488 (residues defining the compact structure of the RBM ridge-loop, Figure 4). In contrast, we observed highly-frequent genetic changes at the hACE2-binding interface affecting the atypical NGV 501-503 sequon (Figure 4). N501Y is the oldest (January 18^th^, 2020 in Spain) and most frequent (63.8%) SARS-CoV-2 mutation detected up to date for the SGP-RBM. It has spread to throughout continents and now accounts for more than 165,500 reported SARS-CoV-2 genomes in the GISAID database (as of March 20^th^, 2021), mainly from the United Kingdom^32,43^. This contrasting finding between both atypical SARS-CoV-2 sequons is in agreement with the well-established knowledge about N-glycans in humans in which a sequon, although necessary, is not a sufficient criterion for glycosylation^23^. Though we identified the atypical N-glycan substrate N501 (Table 1 and Figure 1), we failed in identifying its N-glycan occupancy in any of the SGP-structures reported in the PDB COVID-19/SARS-CoV-2 resources. Moreover, another four mutations at position 501 also destroy the atypical N-glycan sequon (Figure 4) with a much less severe impact on COVID-19 pandemics than the high rates of transmissibility earned by SARS-CoV-2 due to the N501Y, N501T and probably N501S mutations, whose hydroxyl radicals seems to improve the SGP-RBM binding-affinity by stabilizing the salt bridge between D38 and K353 of hACE2 at the *binding-hotspot* 353 (Figure 3B). More worrisome, the mutation N501Y together with E484K, affecting the atypical N-glycan sequons, threaten the protective efficacy of current vaccines^44^ and present new challenges for monoclonal antibody (mAb) therapy, since emergent SARS-CoV-2 variants of major concern (B.1.1.7 and B.1.351) are markedly resistant to neutralization by vaccine sera and convalescent plasma^45–47^. These escape mutations together with the conservative (NGV 481-483) and variable (NGV 501-503) behavior of both atypical N-glycan sequons could indicate their tuning importance in the pandemic, establishing a dangerous pathogenic equilibrium for SARS-CoV-2 in terms of carriage and transmission. Thus, humans are facing one of the most efficient and deadly pathogens.

Unlike replication, transcription, and translation, N-glycosylation in humans is age-dependent and not driven by a template^48,49^. Protein N-glycoforms depends on several human-host parameters such as golgi structure, inflammation, metabolism, glucose availability, and the expression of glycosyltransferases, glycosidases, nucleotide-sugar transporters, and nucleotide-sugar synthetic pathways enzymes^23,50^. N-glycosylation is a complex biochemical process that reflects the consequences of life-style influences and environmental conditions on individuals with different genetic make-up^51^. Moreover, changes in N-glycosylation patterns of plasma proteins have been observed in various aging-associated diseases^50,51^. In this regard, our findings are also able to provide hints on how this novel coronavirus exploits human glycosylation deficiencies, homeostasis disturbance, or complex related diseases targeting the vulnerabilities of a high-risk group of individuals. Those individuals able to atypically glycosylate SARS-CoV-2 impair the efficient binding of its Spike-RBM to the host receptor hACE2, and those with clear deficiencies in these metabolic pathways are more prone to develop severe symptoms of COVID-19 (Figure 5). Thus, depending on the atypical N-glycan antennae displayed on the NGV 481-483 sequon, steric hindrance may come into play at the SGP-hACE2 binding interface, providing the novel coronavirus with a tool for tuning its strong hACE2-binding capacity and viral entry. This age-dependent modulation of the SARS-CoV-2 pathogenic potential will also depend on the individual genetic background underlying the biochemical pathway responsible for the atypical N-glycosylation in humans. In this scenario, humans are masking key non-self coronaviral glycoproteins with their own atypical N-glycans, and this is a plausible molecular explanation for the asymptomatic phenomenon we observed in the pandemic.

**Figure 5.**
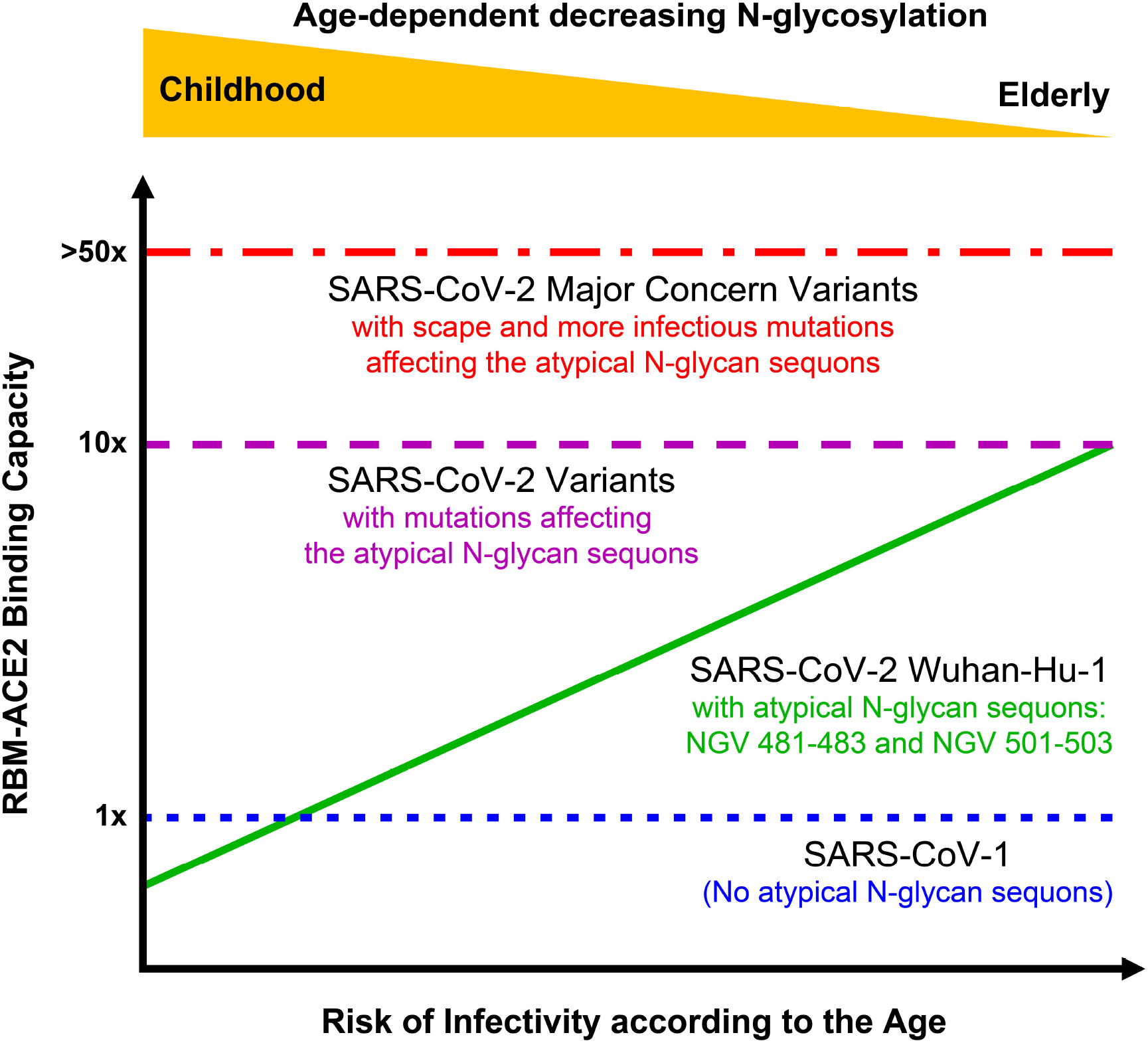
Proposed model of the age-dependent SARS-CoV-2 infection. In contrast to SARS-CoV-1, the SARS-CoV-2 SGP-hACE2 binding capacity can be impaired and/or modulated by the age-dependent N-glycosylation in humans when the atypical N-glycan sequons (NGV 481-483 and NGV 501-503) on the SGP-RBM are used for N-linked glycan attachment. Conversely, humans infected with SARS-CoV-2 variants (B.1.1. 7, B.1.135 and B.1.1.28.P1) making defective the atypical N-glycan sequons are at a higher risk of developing a more severe COVID-19 and dying than are people infected with other circulating variants, regardless of their age and pre-existing health problems.

Finally, despite the contributions that our exhaustive re-analysis of large collections of SARS-CoV-2 genomes and protein structures are making to improve our understanding of the biological behavior of the novel coronavirus, a new window in the molecular biology of this pandemic pathogen is opened. How these atypical N-glycan sequons in the SGP-RBM (differentially glycosylated by humans) could mask immunodominant neutralizing epitopes or how they could affect immunogenicity and lower the effectiveness of the current available vaccines against SARS-CoV-2 are questions we need to answer in the near future. It is difficult to predict the role that NGV 481-483 will play during the massive immunization of human populations, but certainly it has been a determinant in spreading SARS-CoV-2. Thus, it is evident that the atypical N-glycosylation sequons reported in this study for the SARS-CoV-2 RBM has to be considered for future vaccine designs and updates. However, these atypical N-glycan findings for the novel coronavirus need to be more extensively studied and thoroughly to ascertain the implications of their real impact on the transmission and virulence of SARS-CoV-2.

## Materials and Methods

### SARS-CoV-2 reference genome sequence analysis and glycosylation site identification

From the NCBI Virus genome database, we downloaded the available SARS-CoV-2 reference genome sequence (novel coronavirus isolate Wuhan-Hu-1, Accession number: MN908947.3 or NC_045512). By using the SnapGene® software (from GSL Biotech; available at snapgene.com), we manually identified and curated all protein-encoding genes and other relevant traits on SARS-CoV-2 reference genome, according to the updated COVID-19 scientific information publicly available in the literature. Similarly, we performed a genome-wide atypical N-glycosylation site identification in all the SARS-CoV-2 protein-encoding sequences, especially in SGP, according to the sequon motifs NXC and NXV^14,15,26,27^, including the non-functional motifs NPC and NPV (Table 1 and Figure 1). For comparative analysis, we also confirmed the reported prediction of the 22 canonical N-linked glycosylation sites in the SARS-CoV-2 spike-glycoprotein sequence^12,24^, by using the online web-server (http://www.cbs.dtu.dk/services/NetNGlyc), choosing a predication threshold of 0.5.

### Structural visualization and analysis of SARS-CoV-2 spike-glycoproteins

From Protein Data Bank (PDB)^37^ (https://www.rcsb.org/) COVID-19/SARS-CoV-2 Resources (as of March 3^rd^, 2021), we queried and retrieved 276 deposited and publicly available SARS-CoV-2 SGP (172) and RBD (104) structures. Selected entries included 80 crystal X-ray diffraction and 196 cryo-electron microscopy (Cryo-EM) structures. SGP or RBD structures considered for analysis were solved alone (68) or in complexed with their respective hACE2 receptor (28), or bound to antibodies/Fab (153), nanobodies (13), sybodies (7) and/or other ligands/inhibitors-binders (7). Structure visualizations, manual inspection of N-Glycan occupancy and other structural analyses, including residues interactions and structural alignments were performed using the PyMOL program (the PyMOL Molecular Graphics System, Version 2.2.0, Schrödinger, LLC) (https://pymol.org/2/). Spike-glycoprotein structure of SARS-CoV-1 (PDB-ID: 2AJF)^52^ was also downloaded for comparative analysis. All the PDB-ID codes manually classified and inspected are listed in the Supplementary Table S1.

### SARS-CoV-2 spike-glycoprotein alignment and comparison

From the Global Initiative of Sharing All Influenza Data (GISAID) database (https://www.gisaid.org/) we retrieved a total of 730,744 SARS-CoV-2 whole genome sequences (Supplementary Table S2). We used the following criteria to query the database: Host = “Human” AND “Complete Genome Sequence” AND “Low Coverage Exclusion” AND “Complete Collection Date” AND Collection Dates = “December 1^st^, 2019 to February 28^th^, 2021” AND Genomic Sequence Submission = “December 1^st^, 2019 to March 20^th^, 2021”. GISAID considers sequence length >29,000 bp as complete, and further assigns a label of low coverage when the content of unspecified bases (Ns) is >5%. To identify the SGP gene of each SARS-CoV-2 strain, we aligned the complete viral genomes to the SGP gene of SARS-CoV-2 reference sequence (Wuhan-Hu-1, Accession number: MN908947.3 or NC_045512) by using MEGA version X^53^. SGP gene sequences with unspecified bases (N) were filtered out. The selected and aligned SARS-CoV-2 SGP gene sequences (FASTA format) were exported to the SnapGene® software (from GSL Biotech; available at snapgene.com) for translation to amino acid sequences and easy identification of SGP domains, including the RBM and the atypical N-glycosylation sequons, previously detected in the SARS-CoV-2 reference genome. In order to facilitate the screening of SARS-CoV-2 *spike* gene and protein primary sequences, we routinely classified and collected the data from GISAID, according to the transmission course of the pandemic (December 2019 to January 2021) and regions (Asia, Europe, America, Oceania, Africa). Nucleotide and amino acid variations were detected and quantified monthly, aiming to construct a time-lapse Variome of the SGP-RBM in the pandemic. Since SARS-CoV-2 pandemic is a replication-based clonal expansion in a short period of time and SARS-CoV-2 genome evolves mainly by mutation rather than recombination, we used single nucleotide polymorphisms (SNPs) as the main types of genetic variation in SARS-CoV-2 SGP, rather than presence/absence patterns of SARS-CoV-2 encoded genes for analysis of this pathogen. In addition, the SARS-CoV-2 variants of major concern were especially classified and analyzed, according to their appearance, frequency and geographic location.

### Other coronaviruses spike-glycoprotein alignment and comparison

From the NCBI Virus genome database, we downloaded the available SARS-CoV-1, Bat-CoV and Pangolin-CoV reference genome sequences (Supplementary Table 3). By using the SnapGene® software (from GSL Biotech; available at snapgene.com), we identified and aligned all the retrieved SGP sequences of coronaviruses against the SGP gene of SARS-CoV-2 reference sequence (Wuhan-Hu-1, Accession number: MN908947.3). A portion of the SARS-CoV-2 RBM, containing the atypical N-glycosylation sequons was aligned and compared to the corresponding fragment in the sequence of other coronaviruses (21 SARS-CoV-1, 2 Bat-CoVs and 1 Pangolin-CoV).

## Supporting information

PDB Structures

GISAID Genomes

## Data availability

Authors declare that all data and finding supports of this study are publicly available in accessible databases and within this manuscript and its supplementary files.

## Author contributions

*Conceptualization and design of the initial study*: G.G., J.A.H. and C.C.L. *Bioinformatics analysis*: G.G., J.A.H., C.C.L., A.G.M. and C.E.M. *Drafting initial manuscript*: G.G. and A.G.M. *Discussion and interpretation of the findings*: G.G., J.A.H., C.C.L., C.E.M. and S.H. *Review and editing of the manuscript*: G.G., J.A.H., C.C.L., A.G.M., C.E.M. and S.H.

## Acknowledgments

We thank Jessica L. Morales, Luis F. Velez, Laura Muñoz, Santiago Cardona, Andrés F. Castro, Wbeimar Aguilar, Francisco J. Díaz, María T. Rugeles, Andrés Zuluaga, Luis F. García, Gabriel Bedoya and Juan M. Anaya for scientific discussions. We also thank all those who have contributed SARS-CoV-2 protein structures to PDB (https://www.rcsb.org/), genomic sequences to GISAID (https://www.gisaid.org/) and NCBI (https://www.ncbi.nlm.nih.gov/sars-cov-2/) databases, and analyses to Virological.org (http://virological.org/). This study was funded by CELSIA and ISA (Interconexión Eléctrica S.A.) in Colombia and supported by the BFU2017-90030-P grant to J.A.H. from the Spanish Ministry of Science and Innovation. The contents of this manuscript are exclusively the responsibility of the authors and does not necessarily represent the official point of view of funding enterprises and affiliated institutions.

## Competing interests

Authors declare to have no conflict and no competing financial interests.

## Notes

### Competing Interest Statement

The authors have declared no competing interest.

## References

1. WHO. WHO Coronavirus Disease (COVID-19): Dashboard. https://covid19.who.int (2021).

2. Zhou, P. et al. A Pneumonia Outbreak Associated With a New Coronavirus of Probable Bat Origin. Nature vol. 579 https://pubmed.ncbi.nlm.nih.gov/32015507/ (2020).

3. Guan, W. et al. Clinical Characteristics of Coronavirus Disease 2019 in China. N. Engl. J. Med. (2020) doi:10.1056/NEJMoa2002032.

4. Mueller, A. L., McNamara, M. S. & Sinclair, D. A. Why does COVID-19 disproportionately affect older people? Aging 12, 9959–9981 (2020).

5. Cascella, M., Rajnik, M., Cuomo, A., Dulebohn, S. C. & Napoli, R. D. Features, Evaluation and Treatment Coronavirus (COVID-19). StatPearls [Internet] (StatPearls Publishing, 2020).

6. Wan, Y., Shang, J., Graham, R., Baric, R. S. & Li, F. Receptor Recognition by the Novel Coronavirus from Wuhan: an Analysis Based on Decade-Long Structural Studies of SARS Coronavirus. J. Virol. 94, (2020).

7. Wrapp, D. et al. Cryo-EM structure of the 2019-nCoV spike in the prefusion conformation. Science 367, 1260–1263 (2020).

8. Watanabe, Y., Bowden, T. A., Wilson, I. A. & Crispin, M. Exploitation of glycosylation in enveloped virus pathobiology. Biochim. Biophys. Acta BBA - Gen. Subj. 1863, 1480–1497 (2019).

9. Watanabe, Y. et al. Vulnerabilities in coronavirus glycan shields despite extensive glycosylation. Nat. Commun. 11, 2688 (2020).

10. Shajahan, A. et al. Comprehensive characterization of N-and O-glycosylation of SARS-CoV-2 human receptor angiotensin converting enzyme 2. Glycobiology (2020) doi:10.1093/glycob/cwaa101.

11. Grant, O. C., Montgomery, D., Ito, K. & Woods, R. J. Analysis of the SARS-CoV-2 spike protein glycan shield reveals implications for immune recognition. Sci. Rep. 10, 14991 (2020).

12. Watanabe, Y., Allen, J. D., Wrapp, D., McLellan, J. S. & Crispin, M. Site-specific glycan analysis of the SARS-CoV-2 spike. Science 369, 330–333 (2020).

13. Zhao, P. et al. Virus-Receptor Interactions of Glycosylated SARS-CoV-2 Spike and Human ACE2 Receptor. Cell Host Microbe 28, 586 (2020).

14. Sun, S. & Zhang, H. Identification and Validation of Atypical N-Glycosylation Sites. Anal. Chem. 87, 11948–11951 (2015).

15. Sun, S. et al. Site-Specific Profiling of Serum Glycoproteins Using N-Linked Glycan and Glycosite Analysis Revealing Atypical N-Glycosylation Sites on Albumin and α-1B-Glycoprotein. Anal. Chem. 90, 6292–6299 (2018).

16. Xu, W., Wang, M., Yu, D. & Zhang, X. Variations in SARS-CoV-2 Spike Protein Cell Epitopes and Glycosylation Profiles During Global Transmission Course of COVID -19. Front. Immunol. 11, (2020).

17. Andersen, K. G., Rambaut, A., Lipkin, W. I., Holmes, E. C. & Garry, R. F. The proximal origin of SARS-CoV-2. Nat. Med. 26, 450–452 (2020).

18. Lan, J. et al. Structure of the SARS-CoV-2 spike receptor-binding domain bound to the ACE2 receptor. Nature 581, 215–220 (2020).

19. Shang, J. et al. Structural basis of receptor recognition by SARS-CoV-2. Nature 581, 221–224 (2020).

20. Shang, J. et al. Cell entry mechanisms of SARS-CoV-2. Proc. Natl. Acad. Sci. 117, 11727–11734 (2020).

21. Xu, C. et al. Conformational dynamics of SARS-CoV-2 trimeric spike glycoprotein in complex with receptor ACE2 revealed by cryo-EM. http://biorxiv.org/lookup/doi/10.1101/2020.06.30.177097 (2020) doi:10.1101/2020.06.30.177097.

22. Walls, A. et al. Structure, Function, and Antigenicity of the SARS-CoV-2 Spike Glycoprotein. Cell vol. 181 https://pubmed.ncbi.nlm.nih.gov/32155444/?from_single_result=32155444&show_create_notification_links=False (2020).

23. Varki, A. et al. Essentials of Glycobiology. (Cold Spring Harbor Laboratory Press, 2017).

24. Shajahan, A., Supekar, N. T., Gleinich, A. S. & Azadi, P. Deducing the N-and O-glycosylation profile of the spike protein of novel coronavirus SARS-CoV-2. Glycobiology (2020) doi:10.1093/glycob/cwaa042.

25. Chuang, G. et al. Computational prediction of N-linked glycosylation incorporating structural properties and patterns. Bioinforma. Oxf. Engl. 28, 2249–2255 (2012).

26. Lowenthal, M. S., Davis, K. S., Formolo, T., Kilpatrick, L. E. & Phinney, K. W. Identification of novel N-glycosylation sites at non-canonical protein consensus motifs. J. Proteome Res. 15, 2087 (2016).

27. Dang, L. et al. Mapping human N-linked glycoproteins and glycosylation sites using mass spectrometry. TrAC Trends Anal. Chem. 114, 143–150 (2019).

28. Berman, H. M. The Protein Data Bank. Nucleic Acids Res. 28, 235–242 (2000).

29. Liu, L. et al. Potent neutralizing antibodies directed to multiple epitopes on SARS-CoV-2 spike. Nature 1–10 (2020) doi:10.1038/s41586-020-2571-7.

30. Zhou, H. et al. A Novel Bat Coronavirus Closely Related to SARS-CoV-2 Contains Natural Insertions at the S1/S2 Cleavage Site of the Spike Protein. Curr. Biol. 30, 2196–2203.e3 (2020).

31. Hul, V. et al. A novel SARS-CoV-2 related coronavirus in bats from Cambodia. http://biorxiv.org/lookup/doi/10.1101/2021.01.26.428212 (2021) doi:10.1101/2021.01.26.428212.

32. Shu, Y. & McCauley, J. GISAID: Global initiative on sharing all influenza data – from vision to reality. Eurosurveillance 22, (2017).

33. Tegally, H. et al. Emergence of a SARS-CoV-2 variant of concern with mutations in spike glycoprotein. Nature 1–8 (2021) doi:10.1038/s41586-021-03402-9.

34. Sabino, E. C. et al. Resurgence of COVID-19 in Manaus, Brazil, despite high seroprevalence. The Lancet 397, 452–455 (2021).

35. WHO. WHO Coronavirus disease (COVID-19): Vaccines. https://www.who.int/news-room/q-a-detail/coronavirus-disease-(covid-19)-vaccines (2021).

36. Wang, Z. et al. mRNA vaccine-elicited antibodies to SARS-CoV-2 and circulating variants. bioRxiv 2021.01.15.426911 (2021) doi:10.1101/2021.01.15.426911.

37. PDB. RCSB - Protein Data Bank. https://www.rcsb.org/ (2020).

38. Dunn-Walters, D. Effect of somatic hypermutation on potential N-glycosylation sites in human immunoglobulin heavy chain variable regions. Mol. Immunol. 37, 107–113 (2000).

39. Anumula, K. R. Quantitative glycan profiling of normal human plasma derived immunoglobulin and its fragments Fab and Fc. J. Immunol. Methods 382, 167–176 (2012).

40. Boni, M. F. et al. Evolutionary origins of the SARS-CoV-2 sarbecovirus lineage responsible for the COVID-19 pandemic. Nat. Microbiol. 1–10 (2020) doi:10.1038/s41564-020-0771-4.

41. Gussow, A. B. et al. Genomic determinants of pathogenicity in SARS-CoV-2 and other human coronaviruses. Proc. Natl. Acad. Sci. 117, 15193–15199 (2020).

42. Zhang, S. et al. Bat and pangolin coronavirus spike glycoprotein structures provide insights into SARS-CoV-2 evolution. Nat. Commun. 12, 1607 (2021).

43. Rambaut, A. et al. A dynamic nomenclature proposal for SARS-CoV-2 lineages to assist genomic epidemiology. Nat. Microbiol. 5, 1403–1407 (2020).

44. Rubin, R. COVID-19 Vaccines vs Variants—Determining How Much Immunity Is Enough. JAMA (2021) doi:10.1001/jama.2021.3370.

45. Wang, P. et al. Antibody Resistance of SARS-CoV-2 Variants B.1.351 and B.1.1.7. Nature 1–9 (2021) doi:10.1038/s41586-021-03398-2.

46. Collier, D. A. et al. Sensitivity of SARS-CoV-2 B.1.1.7 to mRNA vaccine-elicited antibodies. Nature 1–8 (2021) doi:10.1038/s41586-021-03412-7.

47. Greaney, A. J. et al. Comprehensive mapping of mutations in the SARS-CoV-2 receptor-binding domain that affect recognition by polyclonal human plasma antibodies. Cell Host Microbe 29, 463–476.e6 (2021).

48. Vanhooren, V. et al. N-Glycomic Changes in Serum Proteins During Human Aging. https://home.liebertpub.com/rej https://www.liebertpub.com/doi/abs/10.1089/rej.2007.0556 (2007) doi:10.1089/rej.2007.0556.

49. Parekh R, Roitt I, Isenberg D, Dwek R, & Rademacher T. Age-related galactosylation of the N-linked oligosaccharides of human serum IgG. The Journal of experimental medicine vol. 167 https://pubmed.ncbi.nlm.nih.gov/3367097/ (1988).

50. Vanhooren, V. et al. Protein modification and maintenance systems as biomarkers of ageing. Mech. Ageing Dev. 151, 71–84 (2015).

51. Gudelj, I., Lauc, G. & Pezer, M. Immunoglobulin G glycosylation in aging and diseases. Cell. Immunol. 333, 65–79 (2018).

52. Li, F., Li, W., Farzan, M. & Harrison, S. C. Structure of SARS Coronavirus Spike Receptor-Binding Domain Complexed with Receptor. Science 309, 1864–1868 (2005).

53. Kumar, S., Stecher, G., Li, M., Knyaz, C. & Tamura, K. MEGA X: Molecular Evolutionary Genetics Analysis across Computing Platforms. Mol. Biol. Evol. 35, 1547–1549 (2018).

